# Learning of Active Binocular Vision in a Biomechanical Model of the Oculomotor System

**DOI:** 10.1101/160721

**Authors:** Lukas Klimmasch, Alexander Lelais, Alexander Lichtenstein, Bertram E. Shi, Jochen Triesch

## Abstract

We present a model for the autonomous learning of active binocular vision using a recently developed biomechanical model of the human oculomotor system. The model is formulated in the Active Efficient Coding (AEC) framework, a recent generalization of classic efficient coding theories to active perception. The model simultaneously learns how to efficiently encode binocular images and how to generate accurate vergence eye movements that facilitate efficient encoding of the visual input. In order to resolve the redundancy problem arising from the actuation of the eyes through antagonistic muscle pairs, we consider the metabolic costs associated with eye movements. We show that the model successfully learns to trade off vergence accuracy against the associated metabolic costs, producing high fidelity vergence eye movements obeying Sherrington’s law of reciprocal innervation.

## I. INTRODUCTION

Human infants’ ability to autonomously learn how to make sense of the world around them is truly remarkable. An early milestone in this development is the acquisition of a well-calibrated active binocular vision system capable of detecting and representing binocular disparities and performing accurate vergence eye movements to fixate an object with both eyes. The mechanisms underlying the normal acquisiton of this early developmental hallmark are still poorly understood. We have recently proposed a theoretical framework called Active Efficient Coding (AEC) to describe such learning and self-calibration of sensorimotor loops [1], [2]. In a nutshell, AEC is a generalization of classic theories of efficient coding to active perception. It postulates that biological sensory systems not only strive to encode sensory signals efficiently [3], [4], but they also utilise their own motor behavior, such as their eye movements, to facilitate this objective of efficient encoding of sensory information. Specifically, eye movements that allow the input to be encoded more efficiently are reinforced. Based on this idea a number of self-calibrating models for the development of active binocular vision and active motion vision have recently been developed and validated on robots [5]–[7].

To shed light on the development of binocular vision in human infants, rather than humanoid robots, it is important to carefully consider the complexities of the human oculomotor system. This is particularly important for the study of developmental disorders of binocular vision, which can be rooted in abnormalities of an infant’s oculomotor system [8]–[10]. To address this issue, we have recently developed a detailed biomechanical model of the human oculomotor plant called *OpenEyeSim* [11]. OpenEyeSim combines a sophisticated model of the six extraocular muscles (EOMs) per eye with the simulation and rendering of a virtual environment, which allows us to study closed-loop visually guided behavior.

Here, we utilize the AEC framework and OpenEyeSim to develop the first model of the self-calibration of active binocular vision in a detailed biomechanical model of the human oculomotor system. Since the EOMs activate the eyes in agonist-antagonist pairs, the oculomotor system is *redundant*: Infinitely many combinations of muscle activations will produce the exact same gaze direction. To resolve this redundancy, we incorporate metabolic costs of eye movements, which the model learns to minimize. This results in vergence eye movements obeying Sherrington’s law of reciprocal innervation [12]: as one eye muscle contracts, its antagonistic partner relaxes. We demonstrate that the model is able to self-calibrate the redundant oculomotor system, learning to perform vergence eye movements with sub-pixel accuracy on a time scale consistent with findings on human infants.

## II. METHODS

### Model Overview

The model consists of three distinct stages (see Fig. 1) that are explained in detail in the following sections. At first, the images from left and right eye are pre-processed and dissected into patches. The patches from left and right eye are then combined into binocular patches. These are encoded by a sparse coding model in the form of activations of binocular basis functions. These activations and the activations of the extra-ocular muscles serve as a representation of the current state and are used to generate vergence movements via reinforcement learning. The reinforcement signal is composed of the reconstruction error of the sparse coder as an indicator of the efficiency of encoding and the metabolic costs generated by innervating the eye muscles.

**Fig. 1.**
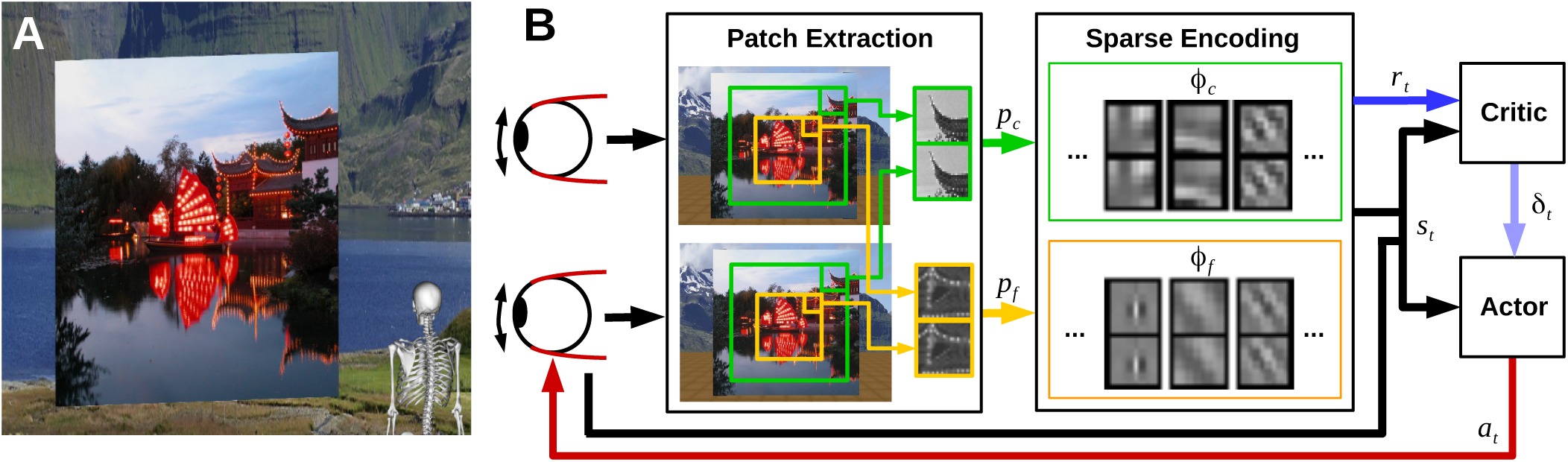
Model overview. **A** The agent looking at the object plane in OpenEyeSim. **B** One image is generated per eye. Binocular patches are extracted in a coarse scale (green boxes) and a fine scale (yellow boxes) with different resolutions. These patches are encoded by activations of basis functions via sparse coding and combined with the muscle activations to generate a state vector (black arrows). While this vector is fed into the reinforcement learning architecture, the sparse coding step also generates a reconstruction error that indicates the efficiency of encoding. We use this signal as reward (blue arrow) to train the critic, which in turn evaluates states to teach the actor (light blue arrow). Finally, the actor generates changes in muscle activations (red arrow), which result in rotations of the eyeballs and a restart of the perception-action cycle.

### Simulation Environment

The action-perception cycle is simulated within the OpenEyeSim platform [11], [13] which contains a detailed biomechanical model of the human extra-ocular eye muscles and renders a virtual environment (VE). This platform is built on top of the OpenSim [14] software for biomechanics simulations. The visual stimulus set used for the experiments was the *man made* section of the McGill Calibrated Color Image Database [15], that consists of outdoor scenes in urban environments. The agent is modeled in the VE by two eyes at a height of 1.7 m with an interocular distance of *d*_*E*_ = 5.6 cm. Two cameras represent eyes which record the scene the agent is looking at. The stimulus is presented on a 3 m *×*3 m plane perpendicular to the gaze direction, which is positioned from 0.5 m to 6 m in front of the agent. Each visual stimulus has a resolution of 1920 px *×*1920 px. The size of the plane was chosen such that for each camera image pixel there is a correspondence of at least 4 px on the stimulus plane to reduce aliasing effects.

As the stimulus position is always vertically and horizontally aligned with the eye level of the agent, it is sufficient to just include the *medial rectus* and the *lateral rectus*, which are rotating the eye around its yaw axis. A landscape scene is used as the background of the VE to prevent the agent from receiving trivial inputs when diverging the eyes from the object plane.

### Image Processing

The first stage of the algorithm starts by rendering the simulated environment from the two virtual cameras corresponding to the left and right eye (each 320 px *×*240 px, covering 50° of visual angle horizontally, with a focal length of *F* = 257.34 px).

Since the use of multiple scales leads to the ability to cope with very different disparity ranges and contributes to the robustness of the system [2], we extract two sub-windows with different resolutions: A coarse scale image, which corresponds to 128 px *×* 128 px in the original image and is down-sampled by a factor of 4 via a Gaussian pyramid, and a fine scale image, which corresponds to 40 px *×* 40 px, without being down-sampled. Patches of size 8 px *×* 8 px from both scales are extracted with a stride of 4 px and normalized to have zero mean and unit norm. Corresponding patches from the left and right image (see Fig. 1) are combined to form 16 px *×* 8 px binocular patches. The previously described parameters were chosen as a trade-off between computational feasibility and biological data concerning the density of photoreceptors and size of ganglion receptive fields on the human retina [16], [17]: Covering a visual angle of 8.8*°* the fine scale corresponds to the central and para-central visual field and the coarse scale with 27.9° to the near peripheral field of view. By down-sampling the coarse scale, we mimic the decreasing resolution towards the periphery.

### Feature Extraction via Sparse Coding

The following step comprises the encoding of the image patches by a sparse coding model. For each scale *S* ∈{*c*, *f*} there is a separate dictionary *ℬ* _*S*_ of binocular basis functions *φ*_*S,i*_, where *|ℬ*_*s*_*|* = 400. There exist *| p*_*f*_ *|* = 81 binocular image patches for the fine scale and *| p*_*c*_ *|* = 49 patches for the coarse scale. A set of 10 binocular basis functions is used to encode each binocular patch *p*_*S,j*_. The *ϕ* _*S,i*_ are chosen by the *matching pursuit* algorithm [18], which tries to yield the best approximation 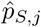 of a patch by a sparse linear combination of basis functions from the respective dictionary:

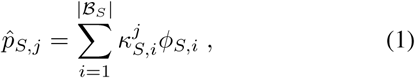

where the vector of *activations 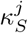* for each patch is only allowed to have 10 non-zero entries.

We calculate the error of this approximation, the *reconstruction error E*_*S*_, as the squared sum of all differences between all patches and their approximations, divided by the total energy in the original patches [2]:

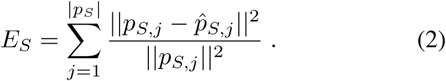

The total reconstruction error *E* is defined as the sum of the reconstruction errors over the two scales: *E* = *E*_*c*_ + *E*_*f*_. The negative of this total reconstruction error serves as a reinforcement signal to train the network that generates the vergence eye movements as described below.

One part of the input to the network controlling the eye movements is a feature vector of size 2 *|ℬ*_*S*_ *|* for both scales combined. Each entry corresponds to the mean squared activation of the *ϕ* _*S,i*_ over all patches. From a biological perspective, the *ϕ* _*S,i*_ correspond to receptive fields of binocular simple cells in visual cortex, while the mean squared activities can be interpreted as the activities of binocular complex cells that pool the activities of simple cells over a larger region of the image.

In our approach, the *ϕ*_*S,i*_ are initialized randomly and adapted during training to represent the input stimulus in the best way through gradient descent on the reconstruction error [19].

### Generation of Eye Movements by Reinforcement Learning

To enable the agent to adjust the fixation plane of the eyes, we simulate the two extra-ocular eye muscles that are responsible for horizontal eye movements: the medial and lateral rectus. OpenEyeSim considers muscle activations in [0, 1] (in arbitrary units).

The model learns muscle commands via reinforcement learning [20]. In particular, we use an actor-critic-framework [21] in a continuous state and action space that implements the idea of positive reinforcement learning named CACLA+VAR [22]. The critic learns to represent the value of the current state *s*_*t*_ described by the state vector. The actor learns to generate motor commands based on the state vector and on the critic’s response.

The critic aims to approximate the *value function*, which is the sum over all (discounted) future rewards starting in the current state: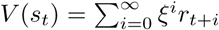where *r*_*t*_, represents the reward achieved at time *t* and *ξ* is the discount factor. We start with random approximations of this value and bootstrap the real solution via *Temporal Difference learning* [23]. The difference between consecutive predictions of the value function, defined by *δ*_*t*_ = *r*_*t*_ + *ξV*_*t*_(*s*_*t*+1_) − *V*_*t*_(*s*_*t*_), is called the temporal difference error (TD-error) and is used to update both value function and policy. Let *θ*^*V*^ represent the parameter vector of the critic’s value function approximation. The update of all entries *i* within this vector follows

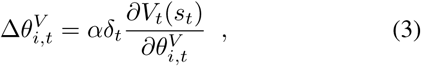

whereby *α* represents the learning rate for updating the critic.

The main characteristic of CACLA+VAR is the update of the actor’s policy network parameter vector *θ*^*A*^. This is only taking place if the last action lead to a better result than expected:

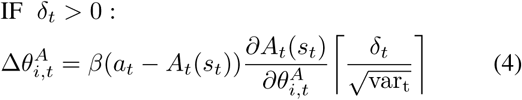

where *β* is the actor’s learning rate, *A*_*t*_(*s*_*t*_) is the action selected by the actor at time *t* and *a*_*t*_ = *A*_*t*_(*s*_*t*_) + 𝒩 (0*, σ*^2^) is the action that is actually executed. This form of action selection is called *Gaussian exploration* and leads to the discovery of new actions by adding zero-mean Gaussian noise to the actor’s output: if the disturbed output leads to a positive TD-error (a positive reinforcement signal), the weights are adjusted to move the actual output closer to the perturbed one. The last term in Eq. 4 scales the update by the magnitude of the TD-error in units of its own standard deviation, which we approximate in an online fashion by 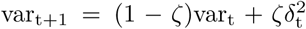 for which we set var_0_ = 1 and *ζ* = 0.001.

For function approximation, we use artificial neural networks. The critic has one input layer and a single output node whose activity represents the value estimate. The actor comprises an input, a hidden, and an output layer. The hidden layer has 50 nodes and uses a hyperbolic tangent activation function. Its output layer has two nodes whose activities are interpreted as changes in muscle activations for the medial and lateral rectus. Since we only learn vergence movements on objects straight in front of the simulated head, the generated muscle activations are symmetric for both eyes.

For both actor and critic, the first layer receives activations from all basis functions from the coarse and the fine scale (as described above), concatenated with the current muscle activations of the medial and lateral rectus. The entries in this state vector *s*_*t*_ are normalized in an online fashion by incorporating Welford’s algorithm [24] to have zero mean and a standard deviation of 0.02.

### Reward Signal

The crucial part in reinforcement learning is the reward function. We distinguish between the following two models: In the first model 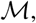 we use the total negative reconstrucstion error as the reward signal. In the second model 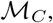 additional metabolic costs resulting from the innervation of the EOMs are incorporated into the reward signal. These are calculated by a muscle model in OpenEyeSim, linearly scaled and subtracted from the total negative reconstruction error to construct the combined reward signal:

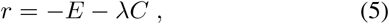

where *E* is the total reconstruction error from the sparse coding step, and *C* are the metabolic costs caused by the current muscle activations at time step *t*. We omit the subscript *t* to increase readability. In 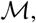 we set *λ* = 0 for training a model without the influence of metabolic costs. For 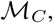 we set *λ* = 0.035, which results in the magnitude of the reward signal being comprised of approximately 20 % of metabolic costs at the start of the experiment.

The calculation of metabolic costs is based on the Umberger model [25] of the human muscle energy expenditure. According to this model, the rate of metabolic energy consumption is calculated as the sum of the activation heat rate of the muscle, the maintenance heat rate, the shortening heat rate, and the mechanical heat rate. For further details the interested reader is referred to [11].

### Experimental Setup

Each experiment comprises a training and several testing phases. The training phase consists of multiple episodes during which a visual stimulus is presented at a certain distance. Each episode consists of 10 iterations or time steps during which the agent tries to accurately fixate the object. Each episode starts with a uniform draw of a stimulus from a set of 100 randomly chosen stimuli from the McGill database [15]. Then the stimulus distance and an initial fixation distance are uniformly drawn from [0.5, 6] m. We initialize the muscles with a pair of muscle activations drawn uniformly at random from the (infinite) set of possible muscle activations of the medial and lateral rectus such that the two eyes verge at the initial fixation distance. For computational reasons, we restrict the muscle activations to [0, 0.2] for the medial rectus and to [0, 0.1] for the lateral rectus within the training and testing phases.

The training phase lasts for 10^6^ iterations (10^5^ episodes) and is intermitted by a total of 13 testing phases. The critic’s learning rate is set to 0.75 and the discount factor to 0.3. We set the learning rate of the actor to 0.5 and let it linearly decline to zero over the course of training. The variance of the Gaussian exploration noise is set to 10^*-*5^.

During a testing phase the learning is turned off and no Gaussian noise is added to the generated muscle activations. A set of 40 randomly chosen stimuli of the McGill database, disjoint from those in training, is used. Each of these stimuli is presented at the distances {0.5, 1, 2*, …,* 6} m. The desired vergence angle corresponding to the drawn fixation distance is calculated by

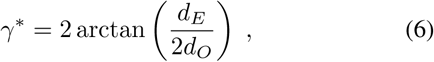

where *d*_*E*_ is the horizontal separation between the eyes and *d*_*O*_ is the object distance. For each distance, initial vergence errors from the set of {–3,–2,–1,–0, …, *η*} deg are applied, where the greatest vergence error *η* depends on the object distance. By doing this, we set the smallest possible initial angle of the eyes to zero which corresponds to the fixation on objects at infinite distance. According to the object distance and the initial vergence error, random muscle activation pairs are drawn. In contrast to the training phase, the agent is given 20 iterations to fixate the object at the testing phase for each scenario to see whether the generated fixations remain stable over a longer period of time. After each iteration, we calculate the vergence error as:

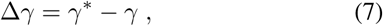

where the current muscle activations determine the current vergence angle *ϒ*.

The assessment of metabolic costs *C* is done in a similar way. For each object distance, there exists one point in the muscle state space where one of the two muscle activations equals zero and the total metabolic cost is minimal. We define the total metabolic cost at this point as *C*^*\**^. *C*_0_ describes the cost at the beginning of every test episode and is determined by the random muscle initializations. The reduction of *C* is calculated by two measures. Δ*C* = *C - C*^*\**^ gives the absolute difference between actual and ideal metabolic costs. Furthermore, we define the relative reduction of metabolic costs by

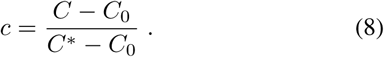

After random initialization, *C* = *C*_0_ and *c* = 0. When the point of optimal metabolic costs is reached, *C* = *C*^*\**^ and *c* = 1. Negative values of Δ*C* indicate an *overshoot*. The system reduces metabolic costs below their optimum value, which it can only do by sacrificing some of its vergence precision.

## III. RESULTS

For each model, the respective experiment was repeated 5 times. As measures of performance, we consider the median and the interquartile range of the absolute vergence error and the median of Δ*C* for all experiments over all 40 test stimuli, all object distances and all initial start vergence errors after 20 iterations as shown in Fig. 2.

**Fig. 2.**
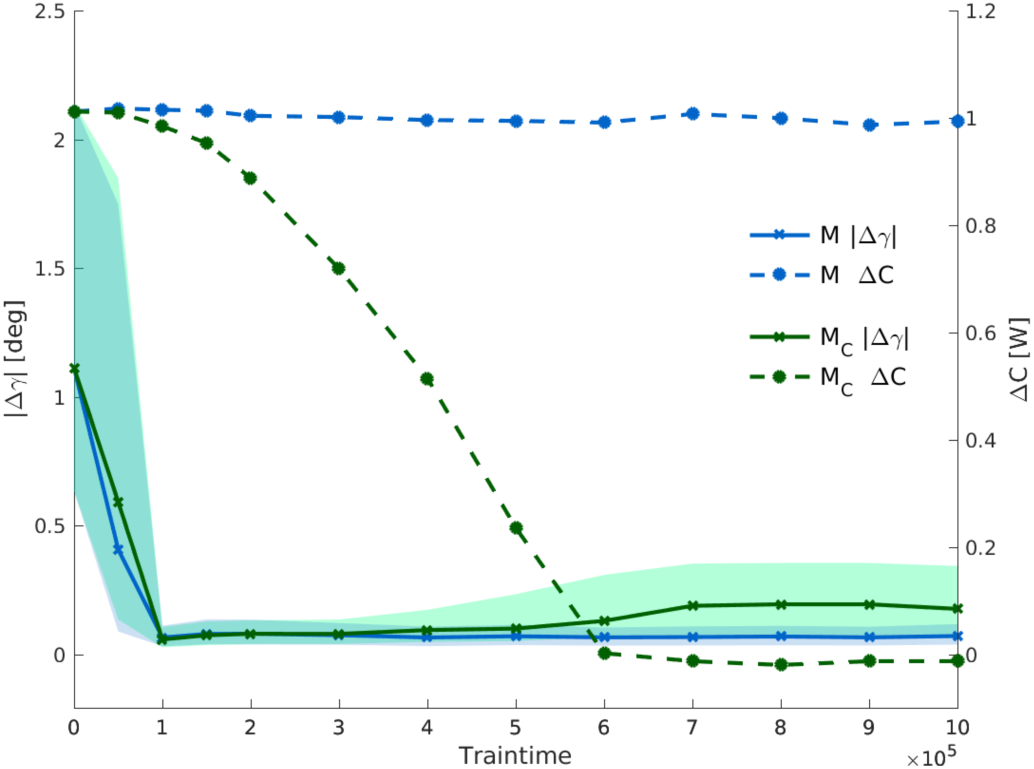
Test performance over the course of training for models with (green lines) and without metabolic costs (blue lines). The solid lines correspond to the median of the absolute vergence error *|* Δ*ϒ|* and the shaded areas to the respective interquartile ranges in degrees. Dashed lines represent the median of the absolute difference of metabolic costs Δ*C* in Watts.

Starting from initially random behavior, both models quickly learn to fixate the given objects as indicated by the decrease in vergence error. This behavior is driven by the reconstruction error generated by the sparse coder. During exposure to binocular images during training, the models learn that the reconstruction error can effectively be reduced, if the images in both eyes are similar [1]. Since actions that reduce the reconstruction error are reinforced, an action policy arises that seeks to generate similar inputs for the left and right eye, that is, reducing disparities by converging the eyes.

However, only M_*C*_ learns to use less muscle activity to fixate the test stimuli. In fact, we can observe a trade-off in the behavior of M_*C*_. The model sacrifices a little bit of accuracy in its vergence eye movements to reduce the metabolic costs below the optimal costs *C*^*\**^ for the perfect vergence angle as discussed in the following.

At the end of training we analyze the change of the performance measures during individual episodes for both models as shown in Fig. 3. Both models greatly reduce *|*Δ*ϒ|* by the second iteration. However, in contrast to M, for M_*C*_ the vergence error slightly increases towards the end of an episode.

**Fig. 3.**
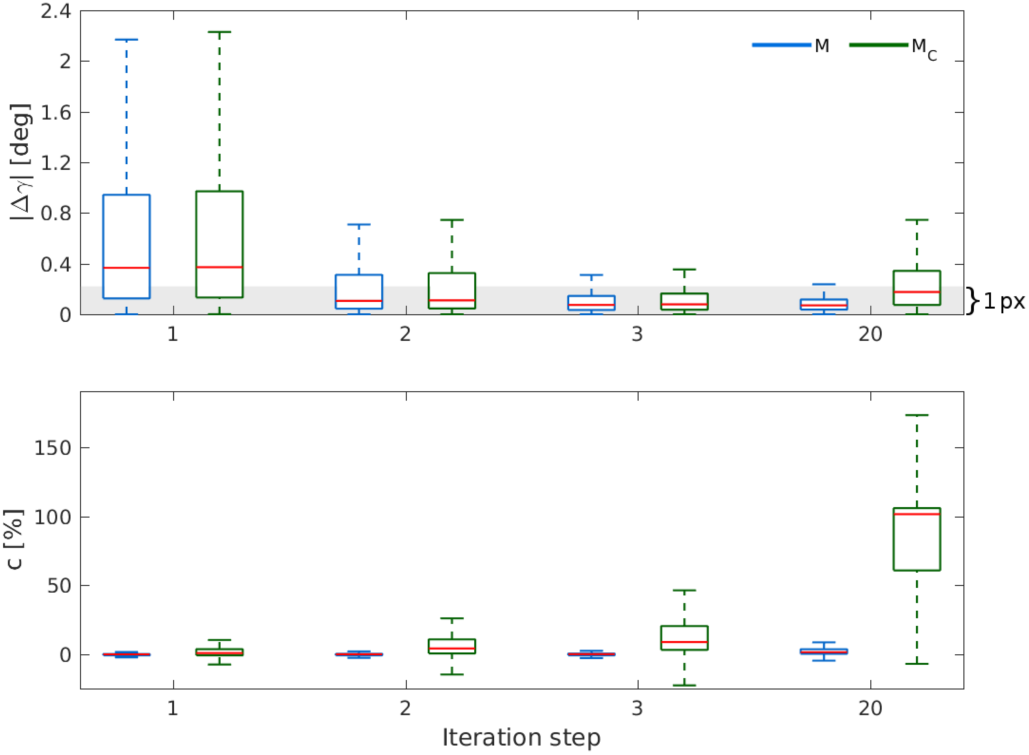
Test performance at the end of training. For the two models with different reward compositions, we plot the reduction of the vergence error and of the metabolic costs after 10^6^ iterations of training using conventional box plots. Displayed are values that are recorded after the 1st, 2nd, 3rd, and 20th iteration of the testing episodes. The gray band in the upper plot depicts an error of 1 px. *c* was scaled to percent in the lower plot.

In the cameras that we simulate, one pixel corresponds to 0.22*°*. ℳ achieves a median *|*Δ*ϒ |* of 0.07*°*. The fourfold interquartile range is 4IQR = 0.319*°*. These measures indicate that the system’s accuracy is in the sub-pixel range. *M_C_* performs worse in this regard with a median *|*Δ*ϒ|* of 0.176*°* and 4IQR = 1.076*°*. In this case, the model accepts a small vergence error to be metabolically efficient.

The relative metabolic costs *c* (in percent) remain in the single digits for Δ*ϒ* indicating no noteworthy reduction of metabolic costs, whereas Δ*ϒ*_*C*_ reduces the metabolic costs by 101.677 %, which indicates its tendency to sacrifice vergence accuracy for smaller metabolic costs.

### Learning Sherrington’s Law of Reciprocal Innervation

In the state space of muscle activations for the medial and lateral rectus different innervations result in the same position of the eye ball, i.e., the same vergence angle and fixation distance. Besides being inefficient in terms of metabolic costs, high muscle activations could also lead to inaccuracies in the visual input due to muscle tremor caused by signal dependent noise [26], [27].

To illustrate this issue of redundancy, we plot example trajectories of muscle activations for the two muscles that are responsible for generating horizontal vergence eye movements in Fig. 4. Displayed are the fixations of two input stimuli, one at 0.5 m and one at 6 m in solid black lines, surrounded by dashed lines that indicate *±* 1 px error in the camera images. The gray scale gradient in the background indicates the total metabolic costs, which increase with greater muscle activity.

**Fig. 4.**
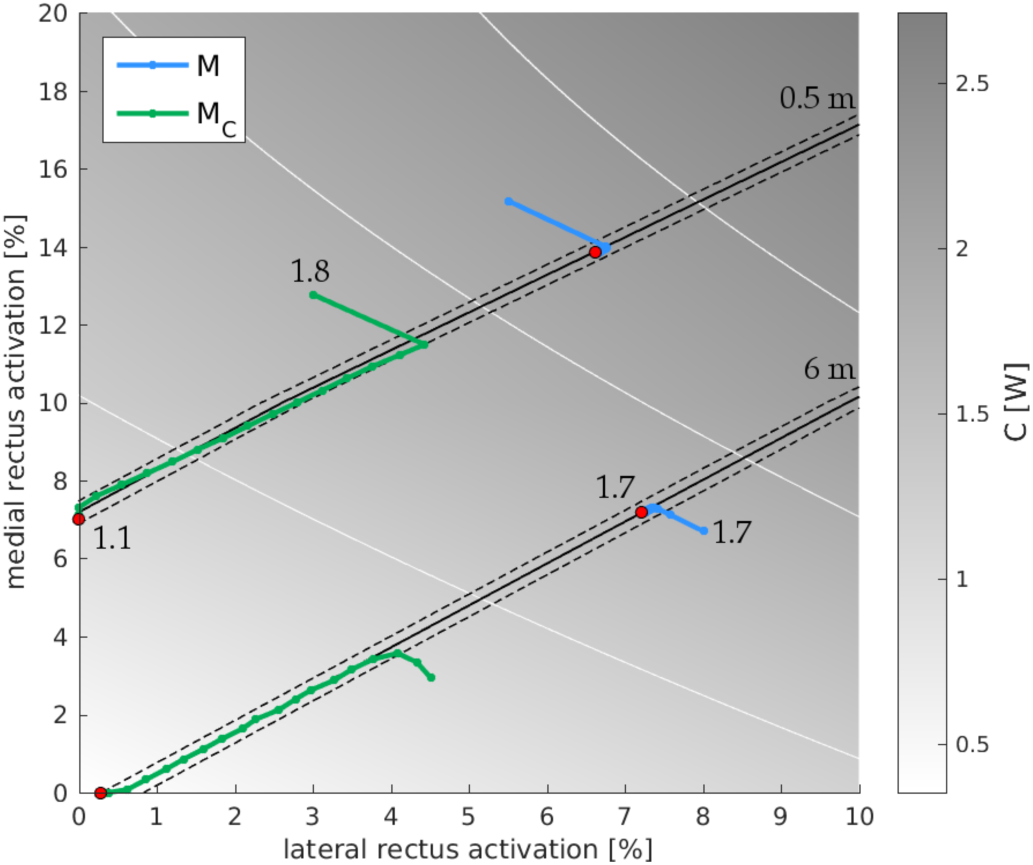
Reduction of metabolic costs in motor space. We plot the resulting metabolic costs in gray values over the activities of the lateral and medial rectus. White lines show contours of constant *C*. Black solid lines indicate muscle activations that result in the same position of the eye and therefore the fixation of the same object depth (here plotted for 0.5 m and 6 m). Red dots mark the end of a trajectory and for two trajectories, the corresponding *C* values are depicted. The model without metabolic costs (M, blue lines) only reduces the vergence error, whereas the model with metabolic costs (M _*C*_, green lines) moves towards a point in motor space that reduces vergence error and metabolic costs at the same time.

For M (blue lines), we observe a straight reduction of the initial vergence error, which is realized by the convergence to the desired vergence angle necessary to perfectly fixate the given stimulus. After approaching *ϒ*^*\**^, M remains in that region of the muscle activation space.

When M_*C*_ is exposed to the same stimuli, it reduces the vergence error at first below *±* 1 px, but then starts to rapidly reduce the costs for muscle activation while still fixating the object. M_*C*_ has acquired Sherrington’s law of reciprocal innervation in that it achieves the correct vergence angle by either relaxing the lateral rectus or the medial rectus as is seen by the green trajectories terminating on the coordinate axes. At the end of the trajectories we observe the model trading off a small vergence error (*≈*1 px) for a further reduction of the metabolic costs by moving slightly away from the line of perfect vergence angle in the direction of the origin of the muscle activation space where metabolic costs are lowest.

## IV. DISCUSSION

We have presented a first model for the autonomous self-calibration of active binocular vision using a detailed biomechanical model of the extra-ocular muscles. Our model extends the Active Efficient Coding framework by introducing metabolic costs in order to resolve the redundancy problem originating from the actuation of the eye ball through agonist/antagonist muscle pairs. We show that the extended model with metabolic costs naturally gives rise to Sherrington’s law of reciprocal innervation. The model learns to choose muscle activations in such a way that when one muscle is activated the other muscle is relaxed.

The our model requires a biologically plausible number of training stimuli if we assume the following: During their first months, infants are awake for around 12 hours per day [28] and fixate objects for less than 5 s [29]. So by the time they have developed their depth perception, which is around 6 months of age [30], they have fixated objects around 10^6^ times which is still an order of magnitude higher than the training episodes required by our model.

In future work, we plan to use the model to investigate possible origins and treatments of developmental disorders of binocular vision such as strabism [31] and amblyopia [32], [33].

## ACKNOWLEDGMENT

This project has received funding from the European Union’s Horizon 2020 research and innovation programme under grant agreement No. 713010 and was supported in part by the German Federal Ministry of Education and Research under grants 01GQ1414 and 01EW1603A, the Hong Kong Research Grants Council under grant 618512, and the Quandt Foundation. L. Klimmasch and A. Lelais are co-first authors.

